# Comprehensive Assessment of Isoform Detection Methods for Third-Generation Sequencing Data

**DOI:** 10.1101/2023.08.03.551905

**Authors:** Yaqi Su, Zhejian Yu, Siqian Jin, Zhipeng Ai, Ruihong Yuan, Xinyi Chen, Ziwei Xue, Yixin Guo, Di Chen, Hongqing Liang, Zuozhu Liu, Wanlu Liu

**Affiliations:** Department of Orthopedic Surgery of the Second Affiliated Hospital, and Centre of Biomedical Systems and Informatics of Zhejiang University-University of Edinburgh Institute (ZJU-UoE Institute), Zhejiang University School of Medicine, Zhejiang University, Hangzhou, Zhejiang 310003, China; Division of Human Reproduction and Developmental Genetics, Women’s Hospital, and Institute of Genetics, Zhejiang University School of Medicine, Zhejiang University, Hangzhou 310006, China; Center for Reproductive Medicine of The Second Affiliated Hospital, and Center for Regeneration and Cell Therapy of Zhejiang University-University of Edinburgh Institute (ZJU-UoE Institute), Zhejiang University School of Medicine, Zhejiang University, Hangzhou, Zhejiang, 310003, China; Zhejiang University-Angel Align Inc. R&D Center for Intelligent Healthcare, Zhejiang University-University of Illinois at Urbana-Champaign Institute (ZJU-UIUC Institute), International Campus, Zhejiang University, Haining, Zhejiang 314400, China; Future Health Laboratory, Innovation Center of Yangtze River Delta, Zhejiang University, Jiaxing, 314100, China; Alibaba-Zhejiang University Joint Research Center of Future Digital Healthcare, Zhejiang University, Hangzhou

## Abstract

The advancement of Third-Generation Sequencing (TGS) techniques has significantly increased the length of sequencing to several kilobases, thereby facilitating the identification of alternative splicing (AS) events and isoform expressions. Recently, numerous computational methods for isoform detection using long-read sequencing data have been developed. However, there is lack of prior comparative studies that systemically evaluates the performance of these software tools, implemented with different algorithms, under various simulations that encompass potential influencing factors. In this study, we conducted a benchmarking analysis of eleven methods implemented in eight computational tools capable of identifying isoform structures from TGS RNA sequencing data. We evaluated their performances using simulated data, which represented diverse sequencing platforms generated by an in-house simulator, as well as experimental data. Our comprehensive results demonstrate the guided mode of StringTie2 and Bambu achieved the best performance in sensitivity and precision, respectively. This study provides valuable guidance for future research on AS analysis and the ongoing improvement of tools for isoform detection using TGS data.

## Introduction

Alternative splicing (AS) is a post-transcriptional regulation mechanism that splices a single kind of pre-mRNA into multiple distinct mature mRNAs, referred to as isoforms. The vast diversity of alternative splicing events significantly contributes to the complexity of the transcriptome and proteome. AS is prevalent in vertebrates, with an estimated 90% of human genes undergoing alternative splicing^1, 2^. Moreover, AS has been observed in invertebrate, fungal, and plant genomes^3^. Accumulated evidence suggests AS plays a crucial role in various biological processes, including cellular differentiation and organismal development, while its dysregulation has been implicated in numerous diseases, including cancer and neurological disorders^4, 5^.

The rapid development of next-generation sequencing (NGS) has revolutionized genome-wide investigations of alternative splicing and isoform characterization. However, AS detection using NGS based RNA-seq relies on recovering splice-junction sites from short NGS read. Consequently, it fails to provide the global picture of full-length transcripts, impeding the study of functional consequences of AS. To overcome this limitation, Third-generation sequencing (TGS) technologies, such as Pacific Biosciences (PacBio) and Oxford Nanopore Technologies (ONT), have emerged^6, 7^. PacBio platform employs single-molecule real-time (SMRT) technology, whereas ONT utilizes nanopores inserted in an electrically resistant membrane^6, 7^. Both platforms enable profiling of full-length RNA transcripts through cDNA (complementary DNA) sequencing, while ONT also allows direct sequencing of native RNA^8–12^. PacBio sequencing includes two modes: 1) continuous long read (CLR), the most common PacBio data type, which yields reads longer than 30 kilobases (kb) but with a higher error rate (8∼15%); 2) High-fidelity (HiFi) sequence reads data type, a circular consensus sequencing (CCS) that generates the highly accurate (>99%) HiFi reads with read lengths of 10∼30 kb^12^. The development and advancement of TGS technologies has significantly increased sequencing read length, enabling precise capture of AS events and isoform structures in TGS RNA-seq data. These advancements open new avenues for comprehensive analysis of transcriptome complexity and functional consequences of AS.

As the advancement of TGS RNA-seq techniques continue, several computational methods have been developed for AS isoform detection from long-read sequencing data. Eight cutting-edge methods, including StringTie2, FLAIR, FLAMES, Freddie, TALON, UNAGI, TAMA, and Bambu, have been published^13–20^ (Table 1). These algorithms can be broadly classified into two categories: guided vs. unguided, depending on whether they require a reference annotation to guide isoform identification. Among the eight software, TALON and FLAMES are guided tools; while Freddie, TAMA, and UNAGI do not require guidance. StringTie2, FLAIR, and Bambu incorporate both guided and unguided modules into their algorithms. To ensure high-confidence isoform-calling, different software implement diverse filtering strategies. For instance, StringTie2 constructs a splice graph for each gene and employs the network flow algorithm for *de novo* transcriptome assembly. Bambu utilizes a machine-learning based probabilistic model for transcript discovery and enables context-aware quantification. FLAIR (guided) and FLAMES perform a step of remapping the raw reads back to the preliminarily assembled transcriptome to filter out potential false positive events based on the abundance of support reads. TALON first labels read with internal priming events, classifies each read as known and novel based on the given reference annotation, and then filters out novel reads with an abundance below a specific threshold. TAMA filters out the low-confidence splice junctions based on their ranking or the amount of mapping mismatch surrounding them. Freddie employs a split-segment-cluster strategy for isoform identification, while UNAGI classifies multiple transcriptional boundaries and filters out splicing events below a specific threshold of their locus coverage. Additionally, most methods perform correction of potential sequencing errors before isoform identification except for FLAIR (unguided) and TALON.

With the increasing number of methods for detecting AS isoforms from TGS data, conducting comprehensive benchmark experiments is crucial to evaluate the applicability of different software under various conditions. However, due to the lack of a ground-truth reference, it is challenging to systemically analyze the accuracy of all eight methods in identifying isoforms across different scenarios. These analyses should consider various potential influencing factors, such as software performance accuracy, the different sequencing platforms, downstream analysis effects, and computational efficiency. Therefore, there is a pressing need for a comprehensive assessment of existing isoform detection methods. Such assessment would not only assist users in selecting the appropriate tools, but also provide valuable guidance to bioinformaticians for further improving existing methods or developing novel approaches for isoform detection.

Using ONT TGS RNA-seq technology along with synthetic, spliced, spike-in RNAs known as “sequins”, previous studies have compared the performance of several isoform detection software^21, 22^. In this study, we systematically compared eight isoform detection software, which correspond to a total of eleven methods, using both simulated and real TGS RNA-seq data. These software were selected based on their relatively complete package documentation and published manuscripts. To generate Nanopore or PacBio TGS RNA-seq data for analyzing the accuracy of different isoform detection software, we developed an upper-level TGS RNA-seq reads simulator called YASIM (Yet Another SIMulator). Numerous TGS Low-Level Read Generators (LLRGs) such as PBSIM1/2/3, NanoSim, and Badread have been published recently^23–27^. YASIM is compatible with several LLRGs such as PBSIM1/2/3, and Badread, and it allows for the simulation of datasets with user-defined read depths, novel AS events, sequencing error rate, reads completeness, and various profiles of sequencing error models. We used the simulated TGS RNA-seq datasets to systematically evaluate the performance of eleven isoform detection methods. Additionally, we assessed the software performance using experimental datasets from various species and cell types collected from previously published data. We also generated Nanopore TGS RNA-seq datasets from naïve and primed human embryonic stem cells (hESCs) in this study. With this paired naïve and primed hESCs TGS RNA-seq dataset, along with our previously published NGS RNA-seq datasets derived from the same conditions, we also performed comprehensive downstream differential isoform usage (DIU) evaluation and experimentally validated one of the DIU isoforms from *RPL39L* (Ribosomal Protein L39 Like) gene using qRT-PCR (Real-Time Quantitative Reverse Transcription PCR)^28^. Furthermore, we assessed the computational performance of the software by evaluating time and memory consumption using an in-house developed profiler.

Overall, our results suggested StringTie2 (guided) and Bambu (guided) as the best AS detection method for TGS data in terms of sensitivity and precision, respectively. FLAIR and FLAMES also exhibited great performance, especially with their additional functional modules for upstream read alignment, downstream differential splicing/expression analysis, and single-cell analysis. In summary, our comprehensive analyses demonstrate the effectiveness of widely used long-read AS detection methods and provide guidance for future research on AS analysis and further development of the tools for isoform detection using TGS data.

## Results

### An Overview of the Benchmark Study

The overall workflow of the benchmark process is illustrated in Figure 1. To address the challenge of lacking a ground truth reference for evaluating the performance of different methods using experimental TGS RNA-seq data, we developed YASIM (Yet Another SIMulator). YASIM facilitates the generation of TGS RNA-seq reads with novel alternative splicing (AS) events. It was specifically designed to support comprehensive benchmarking by allowing users to specify parameters, including read depth, transcriptome complexity index representing number of isoforms per gene, sequencing read completeness, sequencing error rates, and reference annotation completeness. YASIM allows the generation of TGS RNA-seq datasets on different Nanopore and PacBio platforms (Figure S1).

**Figure 1.**
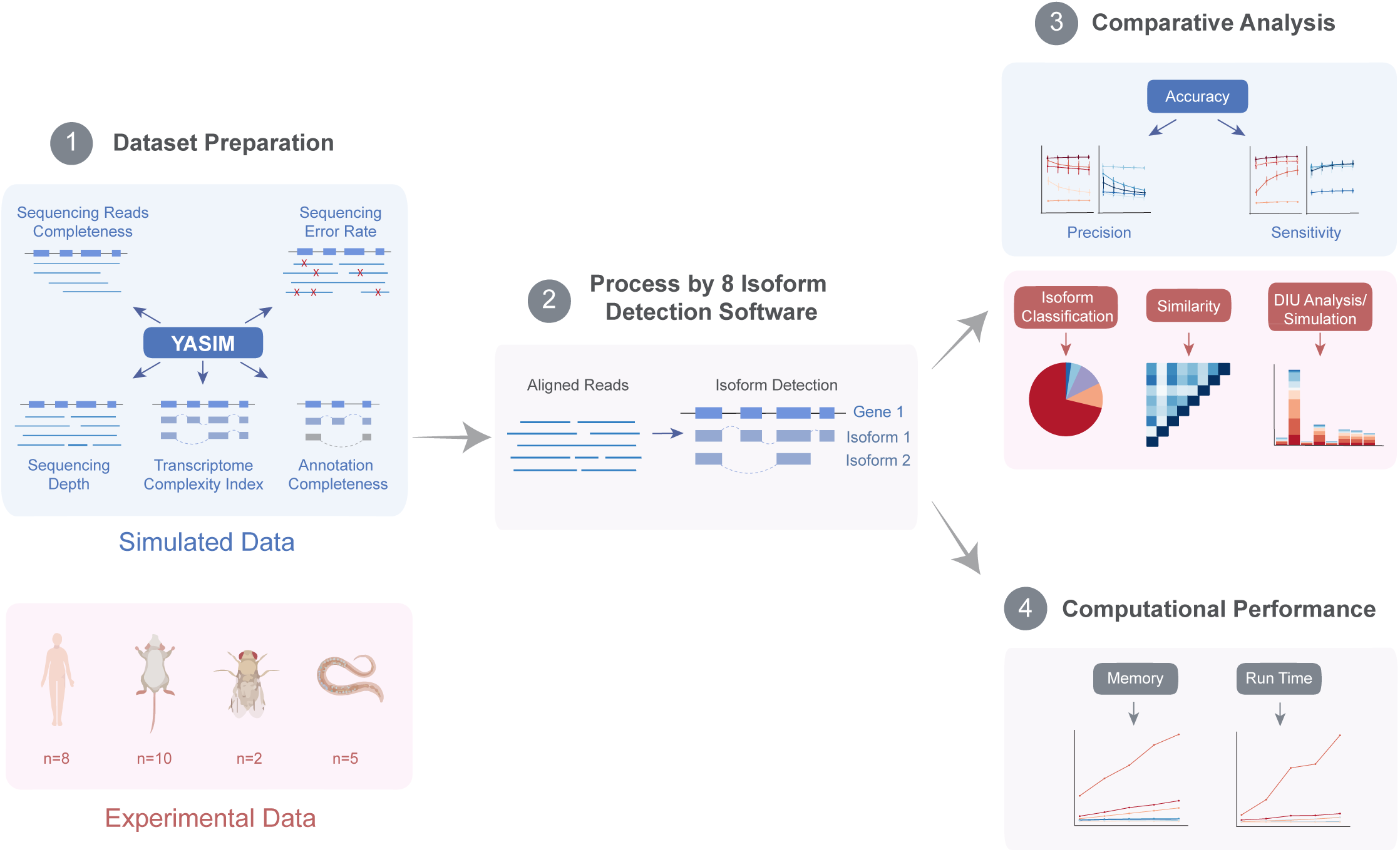
Schematic workflow of the benchmark study. For the preparation of simulated datasets, YASIM was utilized to simulate TGS RNA-seq datasets with variations in sequencing depth, transcriptome complexity index, sequencing read completeness, sequencing error rates, and completeness of the reference annotation. Experimental datasets were obtained from publicly available TGS RNA-seq datasets of four species, including human, mouse, drosophila, and C. elegans. Additionally, in-house TGS RNA-seq datasets were generated from human embryonic stem cells in both Naïve and Primed conditions. The performance of the software was assessed from multiple perspectives, including accuracy, classification of identified isoforms, pairwise similarity between the results, and downstream analysis of differential isoform usage (DIU). Furthermore, computational resource consumption by each method was analyzed.

In the data preparation stage, we utilized YASIM to generate simulated TGS RNA-seq raw data under various conditions, using the *Caenorhabditis elegans* genome as a reference. This approach allowed us to assess the precision and accuracy of different methods. Additionally, publicly available TGS RNA-seq datasets from various species were collected for validation. To enable direct comparison in downstream analysis, TGS RNA-seq data from naïve and primed hESCs were generated using the Nanopore cDNA strategy.

Both the simulated and real data were then aligned to the reference genome using minimap2^29^ with platform-specific parameters. Isoform detection was then performed using different methods. In the comparative analysis, the results obtained from simulated data were compared against the ground truth. Precision and sensitivity values were calculated using GffCompare^30^ to quantitatively evaluate the accuracy of different methods. For experimental data, the detected isoforms from each method were compared against the known reference annotation and classified into different categories of isoforms using SQANTI3^31^. Furthermore, a similarity analysis was conducted by comparing the isoforms detected by different methods using the Jaccard algorithm implemented in BEDtools^32^.

Additionally, downstream differential isoform usage (DIU) analysis was performed using both simulated and experimental data to assess the accuracy and consistency of different methods. Finally, we developed an in-house profiler to evaluate the computational performance, including the speed and memory requirements, of each method across different scales of datasets.

### Generation of Simulated Data with YASIM

To comprehensively assess the software performance, we conducted the benchmarking analysis under various influencing factors. Specifically, we generated five simulated scenarios, each representing different combinations of sequencing depth, transcriptome complexity, sequencing read completeness, sequencing error rate, and reference annotation completeness. Our study encompassed six error models representing both Nanopore and PacBio sequencing technologies: Nanopore R103, reflecting the R10.3 sequencing chemistry released by Nanopore in 2020; Nanopore R94, representing the R9.4 nanopore chemistry developed in 2016; PacBio SEQUEL and PacBio RSII, capturing the error profiles of the PacBio Sequel System released in 2015 and the PacBio RS II System developed in 2013, respectively; and PacBio CCS (Circular Consensus Sequencing) and CLR (Continuous Long Read), representing the two distinct types of PacBio sequencing reads. To generate the simulated data representing these technologies, we utilized the TGS read generator, PBSIM2 and PBSIM3, embedded within the YASIM framework^24, 25^.

For the generation of simulated data used for isoform detection, we generated three replicates of simulated datasets with the six error models under the five different simulation scenarios. We varied the sequencing depth for expressed isoforms, generating depths of 10X, 25X, 40X, 55X, and 70X. Quality control analysis of the sequencing depth for simulated data shows mostly the expected depth for simulated data across all six error models (Figure S2A). For isoform per genes, we generated simulated data with different targeted transcriptome complexity indexes, that positively correlated with the average genome-wide isoforms per genes (Figure S2B). We simulated reads with different levels of read completeness, including full length, 10% or 20% truncation from both 3’ and 5’ end, 10% or 20% truncation of 3’ end, and 10% or 20% truncation from 5’ ends of the TGS reads. The read completeness distribution for the simulated TGS RNA-seq reads distribution under the six error models displayed expected patterns (Figure S2C). We also generated simulated TGS RNA-seq datasets with various sequencing error rates (0%, 5%, 10%, 15%, 20%), and different levels of reference annotation completeness (20%, 40%, 60%, 80%), obtaining simulated data with the expected features (Figure S2D, E). Additionally, we controlled the read length for the simulated datasets. Overall, the read lengths for the simulated datasets were comparable under different simulation scenarios, except for simulated data with truncated read completeness (Figure S3).

### Analyzing Isoform Detection Accuracy Using Simulated Data

We benchmarked a total of eleven modes from the above-mentioned eight software, including those guided by a reference annotation (StringTie2 (guided), FLAIR (guided), FLAMES3, FLAMES10, TALON, Bambu (guided)) and those independent of guidance (StringTie2 (unguided), FLAIR (unguided), TAMA, UNAGI, Bambu (unguided), Freddie). During our evaluation of FLAMES, we discovered that a key parameter, “min_sup_cnt”, used to filter out transcripts with few supports read counts, greatly impacts its performance. Since the bulk RNA-seq module of FLAMES does not provide a default set of parameters, we adopted the configuration file used for running a provided test dataset and modified the threshold of support reads from 10 to 3, which aligns with the default threshold of a similar parameter in FLAIR. The results obtained with number of support reads set to 3 or 10 were both included in our results and were denoted as “FLAMES3” and “FLAMES10”, respectively.

When testing software performance with variations in sequencing depth, Bambu (guided) consistently achieved the highest precision across different depths. Some methods, such as FLAIR (unguided) and TALON, displayed a decrease in precision as sequencing depth increased, particularly on Nanopore datasets (Figure 2A; Figure S4A). In terms of sensitivity, StringTie2 (guided) demonstrated the highest performance on Nanopore datasets, and overall, all methods showed improved sensitivity with increasing read depth (Figure 2B; S4B). Similar trends were observed for PacBio CLR datasets, where changes in read depth had a similar impact on all methods as seen in the Nanopore data (Figure 2A, B; S2A, B). For PacBio CCS datasets, it appeared that the precision of the software were not influenced by the changes in read depths, except for TALON, which exhibited a decrease in precision as sequencing depth increases. However, this decreasing trend was less significant compared to the results from CLR and Nanopore data (Figure 2A; S7A). TAMA achieved the highest sensitivity across datasets with different read depths, and the sensitivity values of all methods also increased as sequencing coverage increased. However, the increasing trend was less remarkable compared to the CLR and Nanopore data, while FLAMES3 was more influenced by the changes in sequencing depth (Figure 2B; S4B).

**Figure 2.**
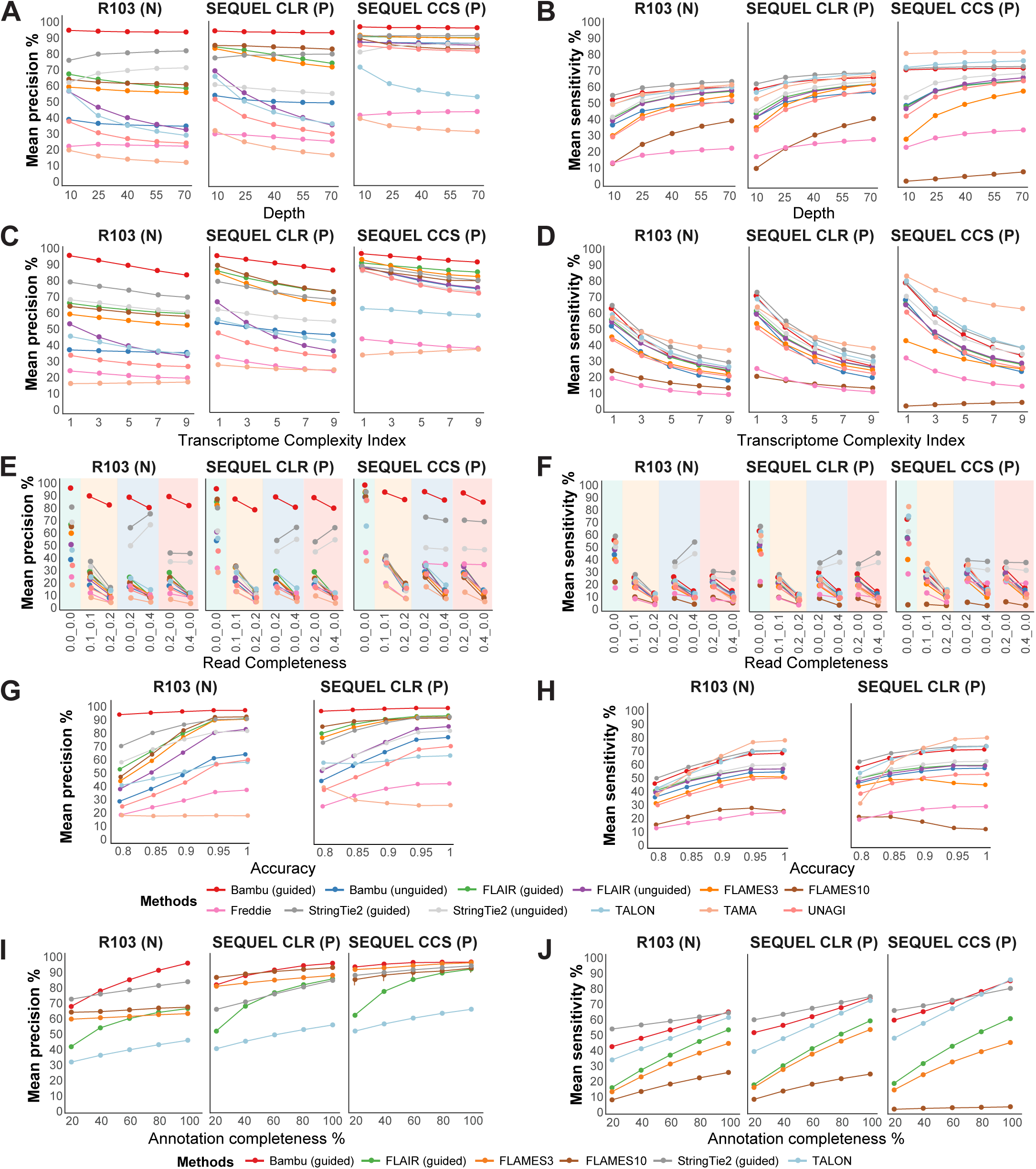
Accuracy of software performance on simulated datasets of Nanopore R103, Pacbio SEQUEL CLR, and CCS. **A, B.** Precision (A) and sensitivity (B) of the performance of tested methods on simulated data of Nanopore R103, Pacbio SEQUEL CLR, and CCS with different read depths (10X, 25X, 40X, 55X, 70X with three replicates for each sequencing platform, n=45 in total). **C, D.** Precision (C) and sensitivity (D) of the performance of tested methods on simulated data of Nanopore R103, Pacbio SEQUEL CLR, and CCS with different transcriptome complexity index (transcriptome complexity index equal to 1, 2, 3, 4, 5, with three replicates for each sequencing platform, n=45 in total). **E, F.** Precision (E) and sensitivity (F) of the performance of tested methods obtained on simulated data of Nanopore R103, Pacbio SEQUEL CLR, and CCS with different read completeness (0.0_0.0: 100% complete, 0.1_0.1: 10% truncated from both ends; 0.2_0.2: 20% truncated from both ends; 0.2_0.0: 20% truncated from 5’ end; 0.4_0.0: 40% truncated from 5’ end; 0.0_0.2: 20% truncated from 3’ end; 0.0_0.4: 40% truncated from 3’ end, three replicates for each sequencing platform, n=63 in total). **G, H.** Precision (G) and sensitivity (H) of the performance of tested methods on simulated data of Nanopore R103, Pacbio SEQUEL CLR, and CCS with different read accuracy (0.8, 0.85, 0.9, 0.95, 1, three replicates for each sequencing platform, n=30 in total). **I, J.** Precision (I) and sensitivity (J) of the performance of tested methods on simulated data of Nanopore R103, Pacbio SEQUEL CLR, and CCS with different annotation completeness (20%, 40%, 60%, 80%, 100%, three replicates for each sequencing platform, n=45 in total). N and P represent datasets generated from the Nanopore and Pacbio platforms, respectively. All values are denoted as mean. Standard deviation (SD) is provided in the Source Data file.

Next, we evaluated the influence variations in the number of isoforms, as represented by the transcriptome complexity index (Figure S2B). For the Nanopore datasets, it is important to note that, except for TAMA and Bambu (unguided), several methods showed a tendency to generate more false positive results as the number of isoforms increased (Figure 2C; S4C). Furthermore, StringTie2 (both modes) detected the highest number of true positives when the transcriptome complexity index was less than three, while TAMA exhibited the highest sensitivity for higher transcriptome complexity in the Nanopore datasets (Figure 2D; S4D). The sensitivity values of all methods experienced a significant decline as the complexity of the transcriptome increased for the Nanopore datasets (Figure 2D; S4D). For PacBio CLR datasets, all methods displayed similar trends for precision as the transcriptome complexity index increased compared to the results from the Nanopore data (Figure 2C; S4C). Similarly, there was no significant difference among various transcriptome complexities compared to that of the Nanopore data in terms of sensitivity (Figure 2D; S4D). In the case of PacBio CCS datasets, the precision of all methods, except for TALON and TAMA, decreased as the number of isoforms per gene increased (Figure 2C; S4C). TAMA also exhibited the highest sensitivity value for datasets with various number of isoforms per gene in the PacBio CCS datasets, while all methods, except FLAMES10, identified a lower proportion of true positives as the transcriptome complexity increased (Figure 2D; S7D).

We also evaluated software performance with variations in read completeness. For the Nanopore datasets, most methods, except for Bambu (guided) and StringTie2 (both modes), exhibited inferior performance with less-complete sequencing reads (Figure 2E, F; S4E, F). Surprisingly, Bambu (guided) displayed relatively stable precision even with truncated Nanopore reads (Figure 2E; S4E). StringTie2 (both modes) on Nanopore datasets showed reduced precision and sensitivity as reads became incomplete from both the 3’ and 5’ ends. However, its performance remained relatively stable when reads were truncated from the 5’ end, and its accuracy even improved noticeably when reads were truncated from the 3’ end (Figure 2E, F; S4E, F). For PacBio CLR datasets, the performance of Bambu (guided) and StringTie2 (both modes) with different read completeness also exhibited different trends compared to the Nanopore datasets. Specifically, Bambu (guided) achieved high and stable precision for truncated reads, while the precision and sensitivity for StringTie2 (both modes) increased as reads-completeness decreased from either 3’ or 5’ ends (Figure 2E, F; S4E, F). Interestingly, for the PacBio CCS datasets, the precision and sensitivity for StringTie2 (both modes) was not significantly impacted by the changes in read completeness from either the 3’ or 5’ truncation, which differed from the CLR and Nanopore data (Figure 2E, F; S4E, F).

TGS RNA-seq simulated data with different sequencing accuracy was also examined. For the Nanopore datasets, Bambu (guided) demonstrated consistent high precision while TAMA consistently exhibited low precision with increased sequencing accuracy. Other methods showed higher precision as sequencing accuracy increased (Figure 2G; S4G). Notably, FLAMES3, FLAMES10, and FLAIR (both modes) are more affected by sequencing accuracy compared to other methods for Nanopore datasets (Figure 2G; S4G). StringTie2 (guided) achieved optimal sensitivity for sequencing accuracy values below 0.9, whereas TAMA exhibited the highest sensitivity for sequencing accuracy values above 0.9 on Nanopore datasets (Figure 2H; S4H).

Increasing read accuracy overall showed a positive impact on software sensitivity, except for FLAMES10, which displayed a declining trend after reaching an accuracy level above 90% (Figure 2H; S4H). For PacBio CLR datasets, similar to the Nanopore datasets, increasing read accuracy had a positive impact on the precision performance of different software, except for Bambu (guided), TALON, and TAMA (Figure 2G; S4G). PacBio CCS data were not included for this analysis, since it can only produce highly accurate reads (>99%).

Since the completeness of the reference annotation may also influence performance for isoform detection, we also investigated the precision and sensitivity of software performance at different levels of annotation completeness. We analyzed software including Bambu (guided), FLAIR (guided), FLAMES3, FLAMS10, StringTie2 (guided), and TALON, which allow the input of reference annotation. For the Nanopore datasets, StringTie2 (guided) demonstrated the highest precision when the completeness of the guidance annotation was below 40%, while Bambu (guided) exhibited the greatest precision otherwise (Figure 2I; S4I). StringTie2 (guided) also achieved the highest sensitivity across different levels of annotation completeness (Figure 2J; S4J). Overall, all methods showed improved performance with increased completeness of the reference annotation, although precision of Bambu (guided) and FLAIR were more influenced by the quality of the input reference, while StringTie2 (guided) displayed less impact on sensitivity compared to other methods (Figure 2I, J; S4I, J). For the PacBio CLR datasets, FLAMES10 detected the fewest false-positive isoforms when the completeness of reference annotation was less than 40%, otherwise Bambu (guided) obtained the highest precision (Figure 2I; S4I). Similar to the results with Nanopore data, all methods on PacBio CLR datasets showed a large improvement in accuracy as the completeness of guidance annotation increases, with FLAIR being the most impacted method (Figure 2I; S4I). The sensitivity results for PacBio CLR datasets with different reference annotation completeness displayed similar results to the Nanopore data. For PacBio CCS datasets, the results obtained from different levels of annotation completeness were not that different from those of CLR and Nanopore data, though it appeared the precision of all methods were less influenced by changes in annotation completeness compared to the other two types of datasets (Figure 2I; S4I). The sensitivity analysis for different levels of annotation completeness were similar to those obtained from Nanopore and CLR data (Figure 2J; S4J).

### Comparative analyses with experimental datasets

To evaluate the software performance on real data, we collected a total of 25 real TGS RNA-seq datasets generated using Nanopore (GridION, MinION, and PromethION) and PacBio (CLR reads, sequenced on RS, Sequel, and Sequel II) platforms. These datasets were obtained from previous publications as well as dataset generated in our own laboratory. They encompassed four different species, namely *Homo sapiens*, *Mus musculus*, *Drosophila melanogaster*, *Caenorhabditis elegans*^33–43^ (see Methods and Table S1).

The quality control analysis of the average Q-Score indicated an overall consistent sequencing quality across the experimental data (Figure S5A). The distribution of read lengths for experimental data exhibited a wide range, spanning from several hundred base pairs to several kilobases (Figure S5B). Additionally, the GC content showed a generally consistent pattern across different samples (Figure S5C). Mapping status for the real data revealed an overall mapping rate of over 75%, although the proportion of primary and secondary alignment events varied among different samples (Figure S6A). Analysis of sequencing errors on experimental data indicated generally low error rates (Figure S6B). Depth analysis demonstrated that most TGS RNA-seq experimental data had depths ranging from 10X to 70X, consistent with our simulated data (Figure S6C). Read completeness analysis revealed that the majority of the experimental data exhibited high read completeness, with the exception of four datasets from mouse samples (sample M5 to M8) (Figure S7).

To evaluate the software performance on experimental data, we included StringTie2 (both modes), Bambu (both modes), FLAIR (both modes), Freddie, FLAMES, and TALON in these analyses. UNAGI and TAMA were not included due to their high computational resource requirements. As the ground truths for the experimental data are not known, we compared the results of different methods by aligning them side-by-side using the most widely used annotation as a reference for each species. The detected isoforms were then classified into five types: full splice match (FSM), incomplete splice match (ISM), novel in catalog (NIC), novel not in catalog (NNC), and intergenic (Figure 3A, B, and Table S2). Mono-exonic transcripts were not included in the classification since they lack splice junctions and may introduce potential false positives.

**Figure 3.**
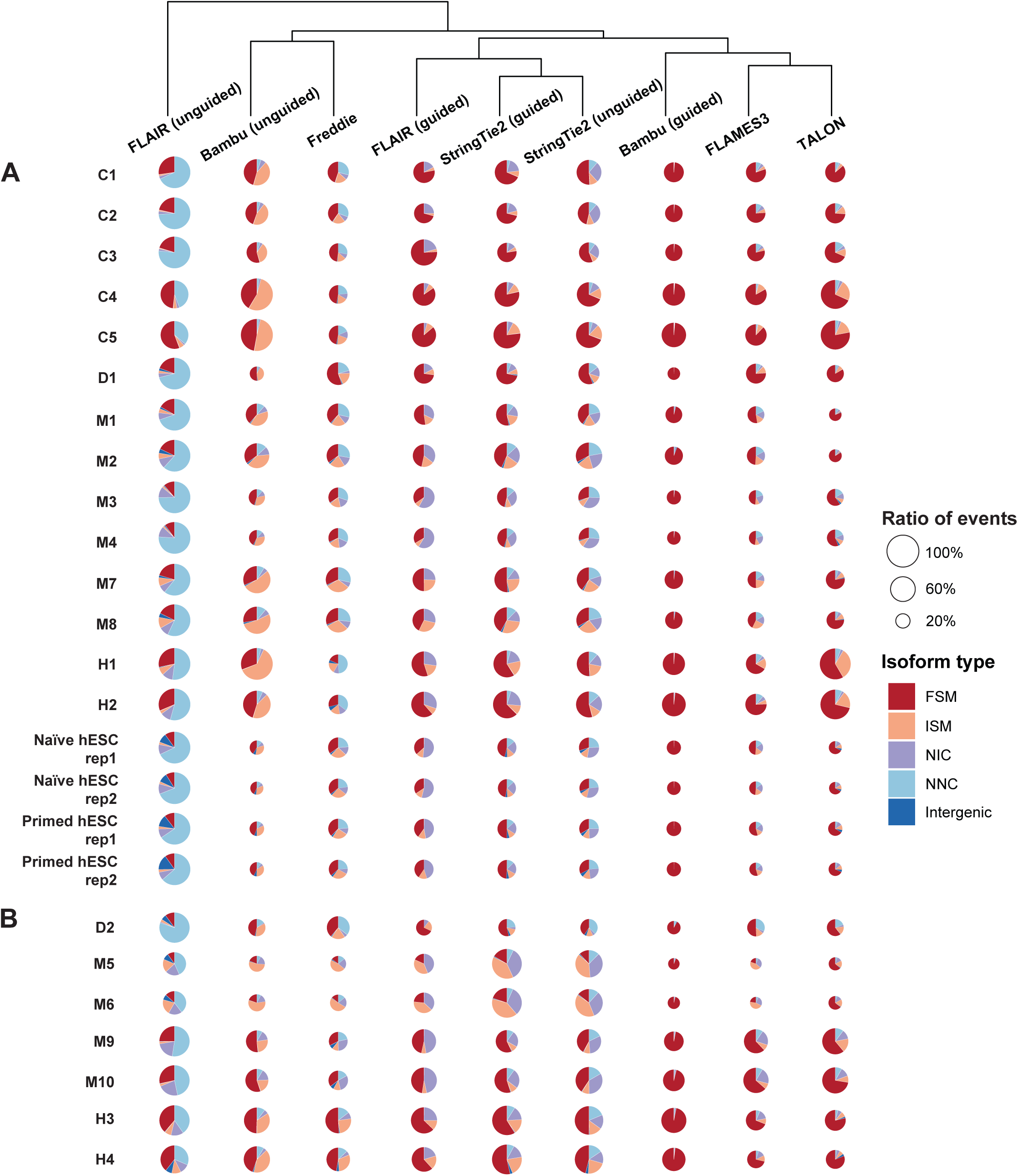
Classification of isoforms detected by different methods from various experimental datasets. **A, B.** Five different isoform types and the total counts of isoform events detected by nine different methods across 25 experimental datasets collected using the Nanopore (A) or Pacbio platform (B). FSM, ISM, NIC, NNC represent full splice match, incomplete splice match, novel in catalog, and novel not in catalog, respectively. The size of each sector is proportional to the largest count of isoform events detected by the nine different methods for each dataset. The results for the nine methods were hierarchically clustered. The publicly available experimental datasets originate from the following sources: C1: L1 larval stage of C. elegans, C2: mix stage of C. elegans, C3: young adult stage of C. elegans; C4: Wildtype C. elegans total RNA replicate 1, C5: Wildtype C. elegans total RNA replicate 2, D1: Drosophila, D2: Drosophila testis, M1: mouse activated CD8 T cell, M2: mouse naïve CD8 T cell, M3: mouse retinal cells (control), M4: mouse retinal cells (glaucomatous), M5: mouse CD4SP cells, M6: mouse CD8SP cells, M7: mouse neural stem cells (E15.5), M8: mouse neural stem cells (P1.5), M9: mouse cerebral cells, M10: mouse hippocampus cells. H1: human Beta cells, H2: human Beta cells treated with cytokines, H3: human Hela cells, H4: human iPSC cells. The TGS RNA-seq dataset on Naïve and Primed hESCs was generated in this study.

When comparing the size of sectors for each method across different datasets, FLAIR (unguided) detected the largest number of isoforms (Figure 3A, B). Bambu (unguided), Freddie, StringTie2 (both) detected a relatively comparable numbers of isoforms. However, Bambu (guided), TALON, and FLAMES may fail to identify certain isoforms detected by other methods. Examining the composition of isoform types detected by different methods in each dataset, guided methods generally detected a higher number of FSM isoforms compared to unguided methods. However, unguided methods were more likely to uncover isoforms that were NIC or NNC, except for Bambu (unguided), which recovered mostly FSM and ISM isoforms. With regards to similarity, StringTie2 (both modes) and FLAIR showed similar patterns between methods, while Bambu (guided), TALON, and FLAMES clustered together. FLAIR (unguided) displayed less similarity with the other methods. Among the methods, Bambu (guided) detected the highest proportion of FSMs, while FLAMES and TALON also showed a high proportion of FSMs. However, Bambu (guided), FLAMES, and TALON all identified a smaller total number of isoforms compared to other methods for most datasets (Figure 3A, B).

In our initial analysis when including the identification of mono-exonic transcripts, some software exhibited a high proportion of NIC and intergenic isoforms, particularly in datasets from mouse and human species. We hypothesized this might be attributed to the high proportion of transposable elements (TE) in the mammalian genome^44^. To validate this hypothesis, we utilized the Nanopore TGS RNA-seq datasets from human naïve and primed embryonic stem cells, which were generated as part of this study. Previous studies have indicated that the dynamic expression of TEs could serve as a hallmark for human naïve and primed hESC^45, 46^, and our earlier research has highlighted their potential functional roles in hESC cell fate determination^28, 47^. Therefore, we utilized this model to conduct an analysis of the isoform types for mono-exonic transcripts identification. Most unguided methods, as well as FLAIR (guided), detected a significant proportion of intergenic isoforms (Figure S8A). Furthermore, the visualization of representative intergenic transcripts specific to naïve hESCs demonstrated a high degree of overlap with previously reported functional TE loci such as LTR5Hs and HERVH/LTR7Y^28, 45, 46^ (Figure S8B, C).

We performed a comparative analysis of the results obtained from different software and quantitatively assessed their similarities using Jaccard statistics, which represent the pairwise overlapping of detected isoforms at the base-pair resolution for each method. Across experimental data from various species and sequencing platforms, Bambu (guided) showed a relatively high level of overlap with TALON, SringTie2 and FLAIR in both guided and unguided modes. Additionally, Bambu (unguided) also exhibited remarkable overlap with StringTie2 (unguided), suggesting a high level of consistency and robustness among these software (Figure 4A, B).

**Figure 4.**
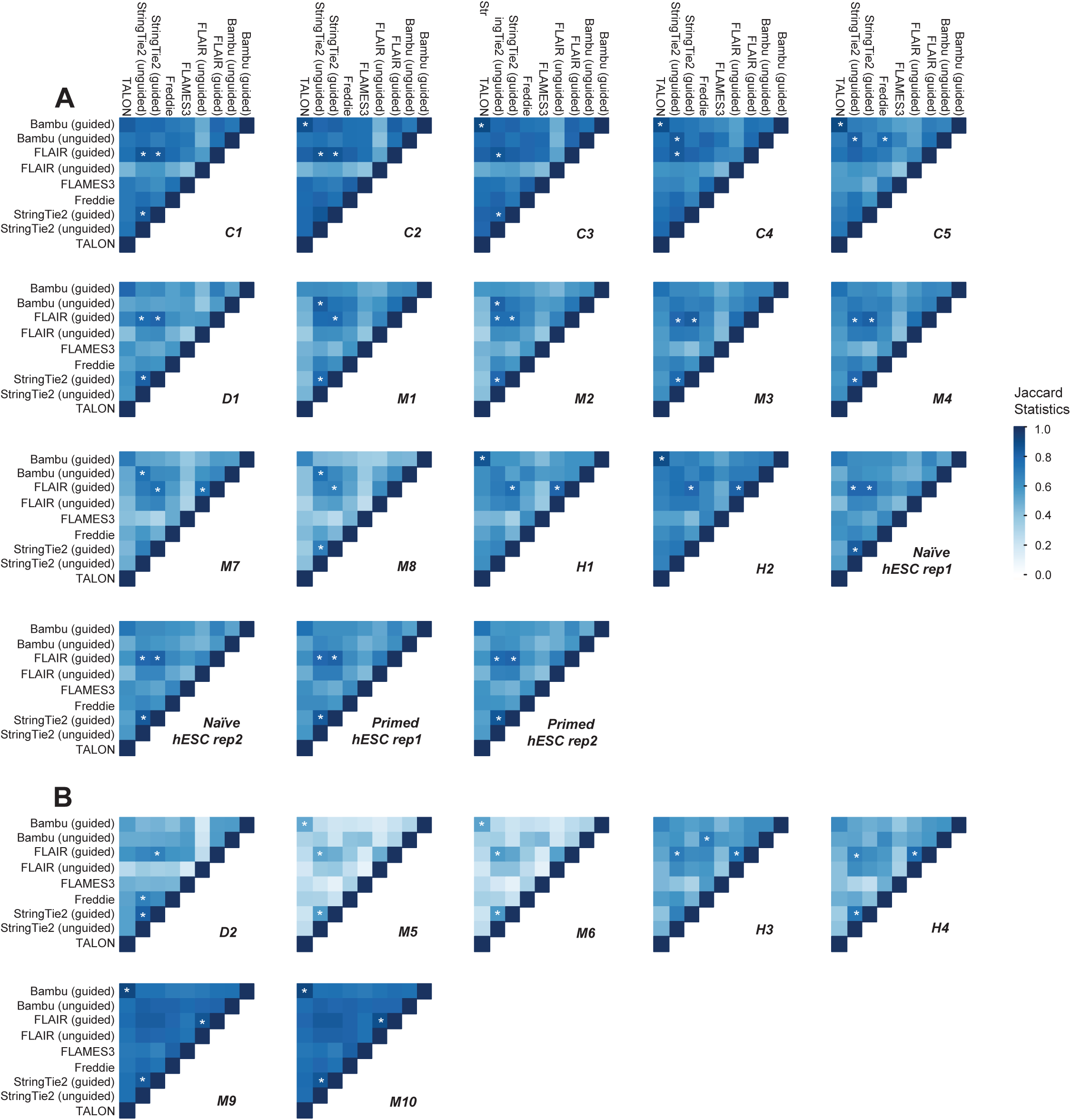
Similarity analyses with experimental datasets. **A, B.** Heatmap showing pairwise Jaccard statistics representing the overlap of isoforms identified by nine different methods across 25 experimental datasets collected using the Nanopore (A) or Pacbio platform (B). The * symbol denotes the top three Jaccard overlaps in each dataset. The publicly available experimental datasets originate from the following sources: C1: L1 larval stage of C. elegans, C2: mix stage of C. elegans, C3: young adult stage of C. elegans; C4: Wildtype C. elegans total RNA replicate 1, C5: Wildtype C. elegans total RNA replicate 2, D1: Drosophila, D2: Drosophila testis, M1: mouse activated CD8 T cell, M2: mouse naïve CD8 T cell, M3: mouse retinal cells (control), M4: mouse retinal cells (glaucomatous), M5: mouse CD4SP cells, M6: mouse CD8SP cells, M7: mouse neural stem cells (E15.5), M8: mouse neural stem cells (P1.5), M9: mouse cerebral cells, M10: mouse hippocam-pus cells. H1: human Beta cells, H2: human Beta cells treated with cytokines, H3: human Hela cells, H4: human iPSC cells. The TGS RNA-seq dataset on Naïve and Primed hESCs was generated in this study.

### Differentially Isoform Usage analyses with both simulated and experimental datasets

Differentially isoform usage (DIU) analysis between groups may usually uncover key condition-related isoforms that may be of potential biological significance. To further investigate the impact of different isoform detection results on downstream analysis, we performed comprehensive DIU analyses on both simulated and experimental TGS RNA-seq datasets from naïve and primed hESCs (See Methods). DIU analysis on the simulated naïve and primed hESCs datasets revealed that those methods which performed best in the previous isoform construction analysis, namely StringTie2 (guided) and Bambu (guided), also demonstrated the highest accuracy in DIU analysis (Figure 5A).

**Figure 5.**
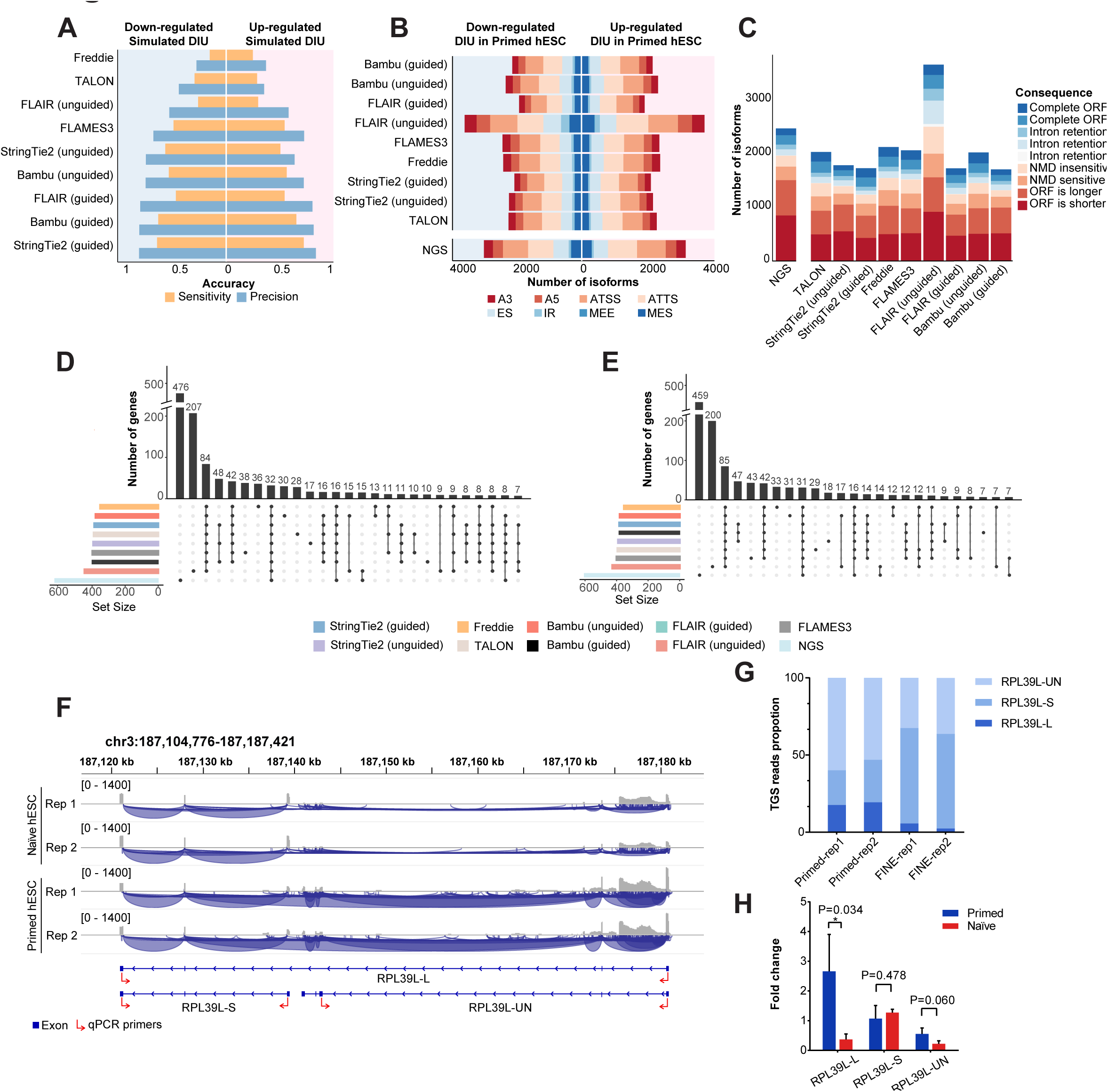
DIU analyses with simulated and experimental data. **A.** Bar plot showing the DIU calling accuracy using simulated data with nine different methods. **B.** Bar plot showing alternative isoform types for up-regulated and down-regulated DIU in Primed hESC compared with Naïve hESCs detected using nine methods. The analysis includes TGS RNA-seq dataset from Naïve and Primed hESC generated in this study, as well as NGS RNA-seq dataset from Naïve and Primed hESC. A3: alternative 3’ splice site, A5: alternative 5’ splice site, ATSS: alternative transcript start site, ATTS: alternative transcript terminated site, ES: exon skipping, IR: intron retention, MEE: mutually exclusive exon, MES: mutually exclusive splicing. **C.** Bar plot showing the number of up-regulated and down-regulated DIU isoforms in Primed hESC with different switching consequences. NMD: nonsense-mediated decay; ORF: open reading frame. **D, E.** UpSet plot showing the number of overlapped DIU genes up-regulated (D) and down-regulated (E) in Primed hESC compared with Naïve hESCs identified by different methods and from the NGS data. **F.** IGV screenshot displaying TGS RNA-seq coverages and splicing junctions of *RPL39L* gene isoforms in Naïve and Primed hESC. The gene model for different transcript isoforms of *RPL39L* is shown under the tracks. *RPL39L-L* and *RPL39L-S* represent isoforms on the gene reference, and *RPL39L-UN* represents the novel isoform detected in this study. Red arrows indicate primers used in RT-qPCR validation experiments. **G.** Bar plot representing the TGS read proportions of the three isoforms of *RPL39L* in Naïve and Primed hESC. **H.** Bar plot representing the RT-qPCR results of the three different isoforms of *RPL39L*. The RT-qPCR was performed using the isoform-specific primers shown in F. Each group involves three biological replicates, and each bar represents the fold change relative to the expression of *RPL39L-S* in Primed hESC. Error bars represent standard deviation. Statistical analysis was performed with a two-sided T-test (\**P* < 0.05).

Additionally, we performed DIU analysis on the naïve and primed hESCs TGS RNA-seq datasets generated in this study. To provide a comparative control, we also included previously published NGS RNA-seq datasets from naïve and primed hESCs cultured under the same conditions. DIU analysis involved applying the isoforms identified by different methods, while the human reference genome was used for the NGS RNA-seq datasets, followed by featureCounts and isoformSwitchAnalyzeR analysis^48, 49^.

The results obtained from the real TGS RNA-seq datasets exhibited variations across all methods, including the number of differentially used isoforms, the distribution of AS event type, and the consequences of isoform switching (Figure 5B, C). Notably, FLAIR (unguided) and the NGS data showed a remarkably higher number of discovered DIUs compared to other approaches (Figure 5B, C). Moreover, the DIU results from guided methods exhibited varying degrees of overlap with each other, while unguided methods, particularly FLAIR (unguided), identified a large number of results that showed no overlap with any other method, possibly due to the higher number of false positive results they generated (Figure 5D, E). The DIU results of NGS data also showed the highest number of unique DIUs compared to results obtained from the TGS datasets (Figure 5D, E).

We further performed experimental validation on *RPL39L*, which is one of the DIUs identified by both Bambu (guided) and StringTie2 (guided) in naïve and primed hESCs (Figure 5F, G). The visualization of TGS RNA-seq tracks for *RPL39L* in naïve and primed hESCs suggested an up-regulation of the RPL39L-Long (*RPL39L-L*) isoform and the presence of a novel isoform structure of *RPL39L*, RPL39L-Unknown (*RPL39L-UN*), in primed hESCs compared to naïve hESCs, while the RPL39L-Short (*RPL39L-S*) isoform may express at a similar level in both conditions (Figure 5F, G). We thus designed isoform-specific qRT-PCR primers and validated the existence of *RPL39L-UN*, as well as the differentially usage for *RPL39L-L* isoform in naïve and primed hESCs (Figure 5H and Table S3).

### Computational performance analyses

We developed a profiler to evaluate the computational efficiency of the benchmarked methods, focusing on two key metrics: total run time and mean memory consumption. Additionally, considering that transcriptome sizes can vary significantly among species, we analyzed the scalability of the software using simulated datasets with different data sizes achieved by varying the sequencing depths. Based on the results, StringTie2 (both modes) demonstrated the fastest speed, highest memory efficiency, and best scalability among the tested methods. FLAMES, FLAIR, and Bambu also exhibited excellent computational performances (Figure 6A, B). However, it is worth noting that some software displayed high time and memory requirements, likely attributed to suboptimal algorithm design and data processing approaches, especially in handling SAM/BAM files (Figure 6A, B).

**Figure 6.**
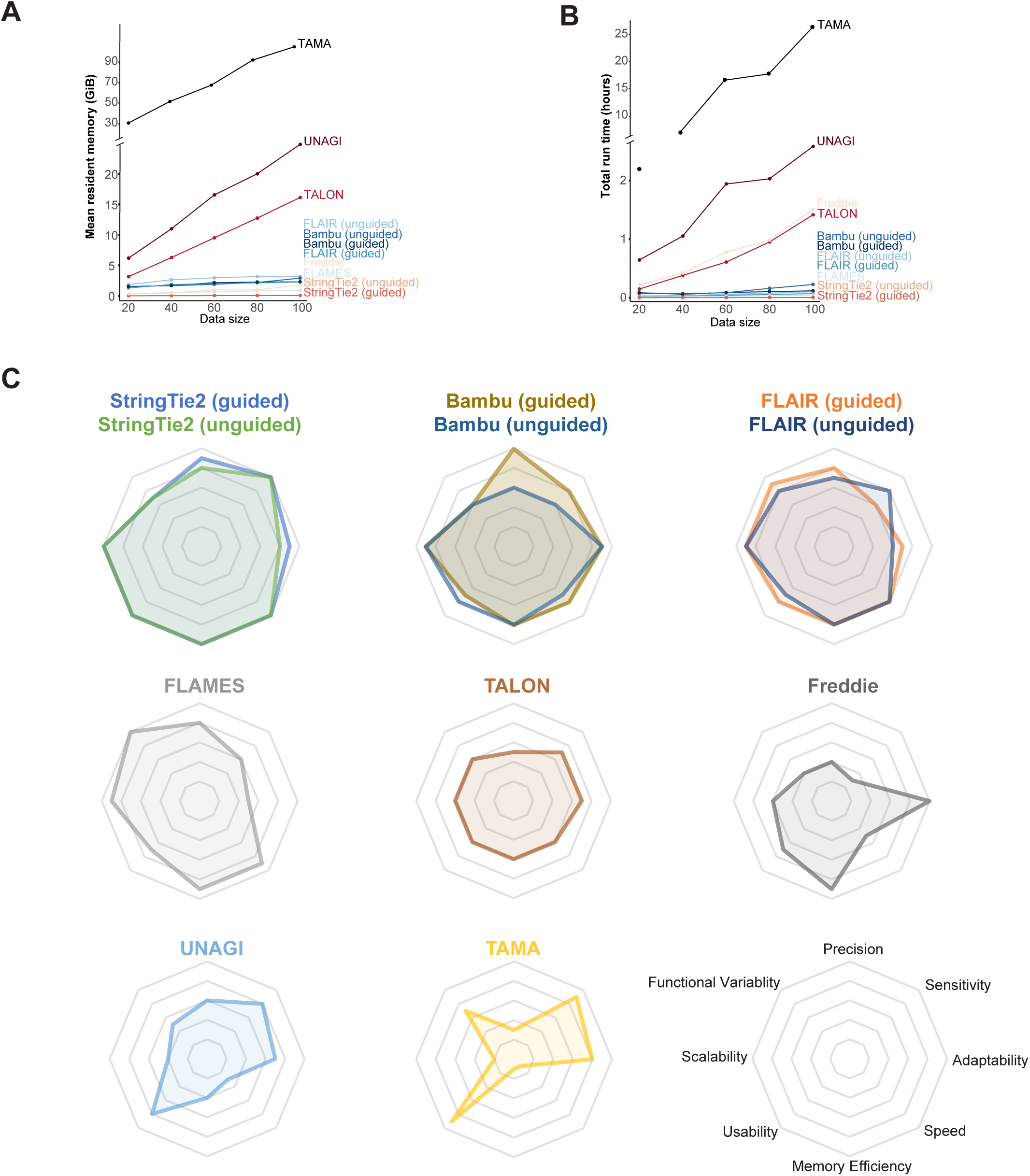
Computational performance analyses and performance summary of the benchmarked methods. **A, B.** Computational performance for mean resident memory consumed (A) and total time spent (B) by each method when processing different scales of data. The data size represents various simulated sequencing depths (20X, 40X, 60X, 80X, 100X) of the datasets used for testing. **C.** Radar charts summarizing our evaluations of eleven different methods across eight aspects, including precision, sensitivity, adaptability, speed, memory efficiency, usability, scalability, and functional variability.

## Discussions

In this study, we conducted a comprehensive analysis of eight computational software tools implemented with eleven different methods for isoform identification in TGS RNA-seq data. We evaluated their performances using a diverse range of simulated and experimental datasets. We also noticed that a new isoform discovery and quantification software, namely ESPRESSO, have recently been developed^50^. Future comprehensive benchmarking on the performance of ESPRESSO would be highly valuable. For the simulation analyses, we selected sequencing depth as a critical factor because this is commonly considered by most software, especially when calculating which isoforms are likely to be false positives and which should be filtered out. Previous studies have indicated a positive correlation between the read coverage and the number of detected AS events, suggesting that reaching certain depths is necessary to detect most isoforms, particularly those with modest abundance^51^. We also considered the number of isoforms per gene as a potential influencing factor to test robustness of software under different data complexities. Genes with a higher number of splice variants pose challenges for accurate reconstruction, as the identification of branch points and systematic analysis of AS events become increasingly difficult as the number of isoforms per gene increases^52^. Additionally, the quality of reference annotation used by certain methods can significantly impact software performance. Inaccurate gene annotations can lead to erroneous isoform identification, whereas more complete annotations are likely to detect a larger proportion of expressed isoforms^53^. To account for the influence of incomplete reads on isoform identification, we simulated different levels of sequencing read completeness in our analysis. Incomplete reads introduce ambiguity in isoform assignment, posing challenges for accurate identification and analysis. We also included sequencing error rate as a factor, considering that long reads, except for CCS reads, tend to have higher error rates compared to short reads (>8-10%). This high error rate presents challenges for alignment and the accurate detection of exon structures in isoforms^13^.

Based on the results, it was observed that increasing the sequencing depths did not evidently improve the precision of the methods. This finding can potentially be attributed to the unique characteristics of TGS data, such as its long read length, which allows a full span across isoforms and relatively even coverage of inter-exonic or intra-exonic regions^12^. It should also be noted that some software tools exhibited an increase in the detection of false positive isoforms as the read depth increased. This phenomenon can be attributed to the relatively high sequencing error rate of TGS. Specifically, software tools that were notably affected by changes in coverage (such as TALON, FLAIR (unguided)) do not incorporate an error correction step prior to isoform detections. Conversely, all methods displayed less sensitivity to changes in read depth when processing highly accurate CCS reads. Furthermore, most methods demonstrated higher precision in CCS datasets compared to other error-prone reads, highlighting the advantage of the high accuracy provided by CCS reads. On the other hand, increasing the sequencing depth positively impacted the sensitivity of the software, enabling the identification of transcripts with relatively low expression levels.

It was observed that as the number of isoforms per gene increased the number of true positives detected by each method decreased. While most methods demonstrated improved performance with fewer erroneous reads, the sensitivity of FLAMES exhibited a declining trend as the sequencing accuracy increased. This can be attributed to one of its key parameters “min_sup_cnt”, which determines the minimum number of aligned reads required for a transcript to be considered. In our analysis, we varied this parameter as 1 (FLAMES1), 3 (FLAMES3) and 10 (FLAMES10) and evaluated its impact on different levels of read accuracy. The precision rates indicated that increasing this parameter may enhance precision under different levels of read accuracy, while FLAMES3 and FLAMES10 demonstrated similar precision rates (Figure S9A). When assessing sensitivity for FLAMES1, FLAMES3, and FLAMES10, we observed that the decreasing trend in sensitivity was less pronounced with a smaller “min_sup_cnt“. Notably, when “min_sup_cnt” was set as 1, the sensitivity value increased as the read accuracy improved, suggesting that error-prone reads that would typically be discarded could be ambiguously assigned to transcripts (Figure S9B). Therefore, it is crucial for users to carefully select the parameter to strike a balance between precision and sensitivity.

The significant negative impact of read completeness on the performance of most software underscores the importance of generating more complete reads for accurate isoform identification. The distinct trends observed in both precision and sensitivity of StringTie2 under single-end truncated reads may be attributed to its usage of splice graph for isoform detection. To further investigate this, we tested the influence of the “-R” parameter in StringTie2, which enables reads cleaning and collapsing without constructing splice graphs. The results revealed that StringTie2 (unguided with “-R”) exhibited poor performance on truncated reads, similar to other methods (Figure S10A, B). Interestingly, the accuracy of StringTie2 (guided with “-R”) during single-end truncation remained relatively unchanged, which is likely due to how StringTie2 refers to the annotated transcripts during its isoform reconstruction process (Figure S10A, B). It is hypothesized that StringTie2 considers a transcript present in the reference as a true positive if the reads overlap with it to a certain percentage and if the transcription start site (TSS) and transcription termination site (TTS) are matched. Additionally, inconsistent trends were observed in the results obtained by StringTie2 from data generated by PBSIM2 and PBSIM3 reads were truncated from the 5’ end. After investigating simulated reads tracks from PBSIM2 and PBSIM3, we hypothesized this discrepancy may be attributed to 5’ degradation in PBSIM2-generated reads, resulting in unmatched TSS and TTS sites, while PBSIM3 did not exhibit this degradation (data not shown). This difference in degradation could potentially influence the process of splice graph traversal and subsequent isoform reconstruction.

It is worth noting that all the software which relies on a known reference annotation demonstrated increased detection for true positives when provided with a higher quality reference annotation. Specifically, methods such as FLAIR and FLAMES3, which utilize the guidance annotation for error correction, appeared to be more susceptible to changes in the quality of reference annotation. This may be attributed to the fact that more true positive reads could be erroneously “corrected” or filtered out when the provided guidance annotation is incomplete.

In the analyses of experimental TGS RNA-seq datasets, the FSM category represents a complete match between detected isoforms and a record in the reference annotation, indicating true positives. FLAIR (guided) and StringTie2 (both modes) exhibited a relatively high proportion of FSMs and displayed similar patterns in terms of the total number of detected isoforms and the distribution of isoform classification compared to other methods. These results are consistent with the findings from that of the simulation analyses, where FLAIR (guided) and StringTie2 (both modes) achieved high precision and sensitivity. Bambu (guided) identified the highest proportion of FSMs, which aligns with the results obtained from simulated data where it demonstrated the highest precision across all simulated scenarios, although its sensitivity may not be as high as that of StringTie2 (guided). However, Freddie and FLAIR (unguided) detected a lower proportion of FSM compared to other methods, consistent with their low sensitivity values in the simulation analyses. For some experimental datasets, FLAMES and Bambu (guided) detected a smaller number of isoforms compared to other methods. Notably, these datasets with a relatively low total number of isoforms detected by FLAMES and Bambu (guided), exhibited a common characteristic of having a higher proportion of non-FSMs types (especially NIC, NNC, and intergenic). The results from all experimental datasets indicate that FLAMES and Bambu (guided) tend to detect fewer of these non-FSMs isoform types, resulting in the recovery of fewer isoforms from this type of data. This observation is consistent with the results of simulation analyses, which showed that FLAMES and Bambu (guided) performed better in precision than in sensitivity. Interestingly, it was observed that certain isoform detection software, such as FLAIR, Bambu, and Freddie can also identify potential TE fragments. The results revealed differences between the TE sequences identified from TGS data and the reference TE annotation, which is predominantly based on short-read sequencing data, particularly at TE boundaries. This suggests the potential improvement of using long-read RNA-seq data for more precise identification of TE structures. The overlaps between isoform detected by different software analyzed at the base-pair level, encompassing FSMs and the novel true positive isoforms. Overall, software with high precision and sensitivity values in the simulation analyses exhibited greater similarity to each other, while tools with relatively lower accuracy demonstrated fewer overlaps with other software.

The diverse range of DIU results obtained from both simulated and experimental data highlights the significant impact of isoform detection on downstream analyses. It is worth noting that a substantial proportion of the DIUs identified from paired NGS data were not detected by any other TGS methods. This could be attributed to the fact that these DIUs were called based on the complete human reference genome annotation during the analysis of NGS datasets, which contains numerous inactive transcripts that may increase ambiguity in read assignment during isoform quantification. Moreover, the nature of NGS data itself, characterized by short read lengths, may increase the likelihood of false positive results. Interestingly, the mouse paralog of the validated DIU *RPL39L* human gene has been proposed to be essential in sperm formation^54^, raising the possibility of further investigating whether these DIU isoforms in naïve and primed hESC have functional implications in the transition between human naïve and primed pluripotency.

Based on the benchmarking results, it can be concluded that among all methods requiring guidance, StringTie2 exhibited the best performance in terms of sensitivity, usability, and computational efficiency, while Bambu (guided) demonstrated the highest precision and relatively high sensitivity (Figure 6C). FLAMES and FLAIR showed lower accuracy but showcased better functional versatility, such as supporting TGS reads mapping, quantification, or single-cell TGS RNA-seq analysis compared to StringTie2 and Bambu (Figure 6C). Among unguided methods, StringTie2 generally outperformed the others in almost every aspect studied. It is worth noting that StringTie2 is the only software that employs a splice graph approach for isoform identification, which represents AS events in a gene as a directed acyclic graph. This approach ensures similar coverage of each exon within an isoform during transcript assembly, thereby avoiding parsimonious yet incorrect results. The splice graph also enables the investigation of AS patterns even under incomplete reference annotation, contributing to the superior performance of StringTie2 in sensitivity and under the unguided mode^55^. Moreover, the use of splice graphs offers computationally efficient by compressing transcriptome data into graph structures, which further contributes to the high computational performance of StringTie2. Bambu stands out with its unique utilization of machine learning models for transcripts discovery, enabling context-specific isoform quantification. It also introduces a precision-focused threshold called the novel discovery rate (NDR), which is calibrated to provide a reproducible maximum false discovery rate across various analyses, thereby avoiding arbitrary per-sample thresholds commonly employed by other isoform detection methods^20^. Our analyses demonstrates that these algorithm designs indeed have the potential to improve the precision of the isoform construction without significantly sacrificing sensitivity for TGS RNA-seq data.

It is important to acknowledge that the benchmarking results obtained from simulated data may not fully capture the complexity of real data, there may also be additional factors influencing the performance of isoform detection tools that were not analyzed in this study, such as GC content^56^. Nevertheless, this benchmark study still offers valuable insights into the comparative effectiveness of most published methods for identifying isoform structures from long-read sequencing data. The findings can serve as guide for the future development of isoform detection algorithms and further investigations into alternative splicing events using TGS data.

## Methods

### Data simulation

We utilized YASIM (version 3.1) to simulate TGS RNA-seq reads containing AS events, which served as the simulated data in this benchmark study. YASIM enables the generation of realistic TGS RNA-Seq with a representative expression profile and AS events, based on distribution models derived from empirical data. YASIM takes reference genome GTF (Gene transfer format) and reference genome FASTA as input and generates corresponding realistic FASTQ sequences as output, along with the ground truth annotation GTF and ground truth expression matrix. The overall workflow of YASIM is as follows: the simulator first selects a specific set of genes as expressed genes and generates new AS events. The number of AS events of each type is derived from empirical data, and the resulting information is written into a GTF file referred as ground truth GTF (gtGTF). The gtGTF is then transcribed to ground truth cDNA according to the reference FASTA. Additionally, an expression profile consisting of transcripts and their corresponding expression levels is generated above the gtGTF. TGS RNA-seq reads are generated by LLRGs according to the error profile that resembles the error profile of a specific sequencer. In this study, we employed PBSIM3 (for PacBio SEQUEL CCS, SEQUEL CLR, RSII CCS, RSII CLR data simulation) and PBSIM2 (for Nanopore R94, R103 data simulation) as LLRGs for simulating PacBio and Nanopore data^24, 25^. The *Caenorhabditis elegans* genome was chosen for simulation, and the ce11 UCSC genome version was used as the reference genome.

Isoform-, Gene- and Sample-Level Depth of simulated and data is calculated and defined as follows. To calculate depths from simulated and real data, the reads are initially unspliced aligned to the reference transcriptome of corresponding species using BWA (NGS data, defaults) or Minimap2 (TGS data, defaults). The depth of each isoform is calculated by dividing the number of primarily aligned bases by the transcribed length of the corresponding isoform. To simplify the calculation process, the depth of each gene is determined by taking the arithmetic mean of the depths of the expressed isoforms within the corresponding gene. Similarly, the depth of each sample is calculated by computing the arithmetic mean of the depths of the expressed isoforms within each sample. The depth of simulated data is calculated using a similar approach, with the exception that, in the simulation, all input isoforms or genes are expressed.

To control the simulated TGS RNA-seq datasets within the dynamic range of gene and isoform expression similar to real data, we applied Gaussian Mixture Model (GMM) and Zipf’s distribution. This allowed us to control the overall RNA abundance, varying it by 10^5^- or 10^6^-fold. In the first step, we generated gene-level depth of each expressed gene. Given targeted mean sequencing depth, this step randomly drew values from a Gaussian Mixture Model (GMM) estimated from several real *Caenorhabditis elegans* TGS RNA-seq and NGS RNA-seq dataset within a specific range (Table S4). By appropriately setting the higher limit, we could generate a distribution with a gene expression variation up to 1000-fold. The second step involved generation of isoform-level depth within each gene. For this, a Zipf’s distribution was assigned as isoform-level expression to each isoform of a gene. The mean of the distribution was equal to the pre-assigned gene-level depth, and the inequality was controlled by parameter “--alpha”. We set “--alpha” to 4, which allowed for a 1000-fold variation among most multi-isoform genes without significantly affecting the means. By applying a similar filtering strategy, we ensured a 1000-fold variation in isoform expression. The third step encompassed the stranded transcription of gtGTF to FASTA, with the isoform name serving as the sequence ID. This was achieved by retrieving stranded exonic sequences from reference genome sequence and concatenating them together. Additionally, this step generated a tab-separated file that recorded statistics for each isoform (e.g., length, GC content, etc.) and a directory where each isoform was stored as separate FASTA. The final step involved the generation of raw reads. This step consisted of two substeps: the generation of reads for each isoform (referred to as “sequencing”, performed by LLRGs) and the assembly of all generated reads into a single file (referred to as “assembling”, performed by an assembler). Firstly, the LLRGs adapter received the isoform sequence, depth, and other customized arguments (e.g., error rate). The adapter then invoked LLRGs, performed cleanup and passed the generated sequence file to the assembler. The assembler reformatted the read ID, performed additional clipping (either from 5’ or 3’ end) if specified, wrote the reads into single file, and recorded statistics such as the actual number of reads generated.

The detailed simulation process in this study was as follows: Firstly, YASIM and *Caenorhabditis elegans* references were installed. AS events were then generated with a transcriptome complexity index set to two as the base gtGTF for all simulations, except for those involving different transcriptome complexity. To simulate TGS RNA-seq data with different number of isoforms per gene, a parameter named transcriptome complexity index (“--complexity” parameter) were applied. To simulate TGS RNA-seq data with varying depths, the “--mu” parameter within YASIM was adjusted accordingly. To simulate TGS RNA-seq data with different read completeness, the “--truncate_ratiio_5p” or “--truncate_ratio_3p” parameters were adjusted to clip a proportion of reads from 5’ end or 3’ end, respectively. For simulating TGS RNA-seq reads with different error rates, the YASIM “--accuracy-mean” parameter, which called the error rate parameter within PBSIM1/2/3, was adjusted. To simulate reference annotation with different completeness, the YASIM “--percent” parameter was applied to randomly discard certain proportion of reference annotation.

All simulated sequencing data was mapped using minimap2 (version 2.17-r941)^29^ with “-ax splice -MD” parameters. The resulting SAM files were sorted, converted to BAM format, and indexed using SAMtools (version 1.15.1)^57^. The sorted BAM files were then processed by StringTie2, FLAMES, FLAIR, Bambu, Freddie, whereas the sorted SAM files were processed by TALON and TAMA. UNAGI, on the other hand, takes FASTQ and the reference genome as inputs, as it is embedded with the alignment process.

For DIU simulation part, YASIM is currently only compatible with UCSC reference genomes. We used the UCSC release of hg38 National Center for Biotechnology Information (NCBI) RefSeq reference annotation to obtain the count matrix for each isoform of naive and primed hESCs TGS RNA-seq datasets. We used featureCounts to generate the count matrix (parameters “-O -L -t transcript -g transcript_id“) (version 2.0.0)^48^. Then, we directly called 96 DIU genes using isoformSwitchAnalyzeR (version 1.8.0) based on the count matrix obtained from featureCounts^49^. The corresponding isoform annotations of these 96 genes were extracted from the reference GTF as ground truth. Additionally, we randomly selected another 100 genes from the reference annotation, excluding the 96 DIU genes, and mixed them into the gtGTF. We then extracted the isoform counts from these 196 genes from the isoform count matrix generated by featureCounts. This resulting isoform count matrix, containing the 196 genes, served as the input for generating expression profiles in YASIM. We generated four corresponding simulated TGS RNA-seq datasets using the Nanopore R103 error model. The simulated reads were then aligned using minimap2 (version 2.17-r941) and processed with the nine benchmarked methods^29^. The assembled transcriptome was extracted from the hg38 reference genome by GffRead (version 0.12.7) based on the isoform annotation provided by each method^30^. Quantification was performed using Salmon (version 1.8.0) with the parameters of “--ont -l U”, and DIU analysis was conducted using isoformSwitchAnalyzeR^49, 58^. To account for any biases introduced by salmon, we also directly called the DIUs using Samlon directly on the gtGTF of the 196 genes. Finally, we compared the DIUs called based on the quantification results obtained from the transcriptome constructed by isoform detection methods with ground truth DIUs to obtain the precision and sensitivity.

### Process of experimental datasets

The Nanopore direct RNA-seq data was aligned with minimap2 (version 2.17-r941) using the parameters “-ax splice -uf -k14 --MD”^29^. The Nanopore cDNA data was aligned using “-ax splice --MD” parameters using minimap2^29^. For the PacBio data, the alignment was performed using “-ax splice:hq -uf --MD” parameters in minimpa2 (version 2.17-r941)^29^. Quality control of NGS naïve and primed hESCs data was performed using FastQC (version 0.11.8) (https://www.bioinformatics.babraham.ac.uk/projects/fastqc/). The first 10 bp of both paired-end reads were trimmed by Cutadapt (version 2.9)^59^. STAR (version 2.7.1e) was used for alignment of NGS RNA-seq data with parameters set as “--outFilterMultimapNmax 1000, --outFilterMismatchNmax 3, --outSAMmultNmax 1”^60^. All resulting SAM files were sorted, converted to BAM, and indexed with SAMtools (version 1.15.1)^57^.

### Cell culture and qRT-PCR validation

Naïve and primed hESCs (H1 cell line, Wicell Research Institute, Inc., WA01-pcbc) were maintained following previously described protocols^28^. Briefly, the primed hESC line H1 were routinely cultured in mTeSR1 medium (Stem cell technologies, 85851) on matrigel (Corning, 354277) pre-coated plate in a normal O2, 5% CO2 incubator. Passaging was performed every 4 days, and the medium was refreshed daily as described before^28^. The naive hESC line was derived from primed H1 line using feeder-independent naive embryonic (FINE) stem cells culture system, as previously described^28, 61^. Adapted H1 cells were cultured in FINE medium under hypoxia conditions (5% O_2_, 5% CO_2_) on plates pre-coated with a reduced growth factor matrigel. Both naïve and primed hESCs were regularly tested for mycoplasma contamination, and all tests showed negative results.

To perform TGS RNA-seq, total RNA was extracted, reverse transcribed, quality controlled, and Nanopore cDNA RNA-seq library was constructed according to the Nanopore Ligation Sequencing Kit 1D (PM) using pore type R9.4.1, and sequenced on PromethION platform by Novogene Co., Ltd.. For the qRT-PCR validation experiment, total RNA was extracted from three independent replicates of naïve and primed hESCs harvested by TRIzol RNA isolation reagents at different passages (Thermo 15596026). The cDNA was synthesized by reverse transcription of 0.5 µg RNA using the RT-PCR kit (Vazyme R222). qPCR analysis was performed using the qPCR SYBR-green kit (Vazyme Q712-03) following the manufacturer’s protocol. Isoform specific primers listed in Table S5 were used to amplify the three *RPL39L* isoforms, with *GAPDH* serving as the internal control gene. The qPCR experiments were performed using the LightCycler® 480 System (Roche).

### Computational performance analyses

The computational performance analyses were conducted on a workstation with an AMD Ryzen Threadripper 3970X 32-Core Processor and 256 GiB 2133MT/S DDR4 memory. The system ran Ubuntu 20.04 LTS with the latest updates and a 5.13.0-44-generic kernel. Memory consumption for each method was measured based on the residential set size, a metric commonly used by other profiler tools. The run time of each software was recorded from the start to the termination of the execution.

The profiler used in this study, proc_profiler, is a process-level profiler that collects metrics such as CPU utilization or memory consumption of processes and child processes in an asynchronous manner. It is implemented in Python (version 3.8) on top of psutil library (version 5.9). The profiler is designed to be executed on GNU/Linux systems only. The workflow of the profiler is as follows. It starts by targeting the command line using the subprocess module of Python and records the process ID (PID) of the targeted process. Then, it generates a dispatcher over this process, which further generates several tracers. Each tracer is a single thread that asynchronously probes various aspects of the process, including CPU usage, memory usage, I/O operations, file descriptor, and current status, using the psutil library. The information collected by the tracers is appended to separate GZipped Tab-Separated Values (TSV) files using appenders and reported to a Command-Line Interface (CLI) frontend. The dispatcher also monitors if the process has spawned new processes (or forked on Microsoft Windows) new processes. If new processes are detected, a new dispatcher is started to monitor these processes. The dispatcher or tracer terminates when the process it is monitoring terminates, and the program exits when the main dispatcher ends. A system dispatcher, which manages tracers that track system-level metrics, is started and terminated at the same time as the main dispatcher. The output of the profiler consists of a folder containing multiple GZipped TSVs files, each capturing different metrics on the monitored process(es).

### More details of the benchmarking process

To ensure the objectivity of the benchmark study, every evaluated method was tested with default parameters. FLAMES does not provide a default set of parameters for its bulk RNA-seq module, so we used the configuration file it included in the software for running a test dataset. We modified the threshold of support read (controlled by the “min_supp_cnt” parameter), from 10 to 3. In other words, the software discarded transcripts with less than 3 aligned reads. This threshold is also the default threshold used by FLAIR, indicated by the “--support” parameter. The results obtained using FLAMES with read support equals to 10 were included in the results and was denoted as “FLAMES10”. For the unguided mode of FLAIR, the correction step was not performed as it requires a reference annotation for guidance. The collapse step was conducted directly without the input of a reference annotation. Results of TAMA and UNAGI were also transformed into GTF before being further analysis. We used GffCompare (version 0.12.6) with default parameters to compare the accuracy of the results obtained with simulated data against ground truths^30^. The transcript level statistics were used as the reference metric. SQANTI3 (4.2) was run in default mode for isoform classification^31^. For the DIU analysis on naïve and primed hESCs datasets, isoforms detected from different samples with each software were first merged together guided by the reference annotation. The merge function in StringTie2 was used with the parameters “--merge -L -G {reference GTF}”, and featureCounts (version 2.0.0) was then used to quantify the isoform expression with “-O -L -t transcript -g transcript_id” parameters based on the merged annotations^48^. For the NGS primed and naïve hESCs data, featureCounts was used to quantify the isoform expression based on the GRCh38.105 reference annotation, using the parameters “-O -t transcript -g transcript_id”. IsoformSwitchAnalyzeR (version 1.8.0)^49^ was used to perform the DIU analysis based on the expression matrix provided by featureCounts. The isoform switching consequence were obtained using the “extractConsequenceSummary()” function, considering selected consequences such as “intron_retention”, “NMD_status”, and “ORF_seq_similarity“. The distribution of AS events for isoforms of differential usage was provided by the “extractSplicingSummary()” function.

Precision is calculated as TP (True Positives) divided by the sum of TP and FP (False Positives)). Sensitivity is calculated as TP divided by the sum of TP and FN (True Negatives)). TP refers to “true positives”, which in this investigation refers to the isoforms detected that match the transcript records in the corresponding ground truth annotation. FN (“false negatives“) are transcripts present in the ground truth but missed in the isoforms identified by the software, while FP refers to “false positives” that are found in the detected isoforms but not recorded by any ground truth. An identified isoform was considered matched with the ground truth (i.e. true positive) if it shared all splice site boundaries exactly compared to an annotated transcript. Regarding the classification of isoforms, FSM refers to the query isoforms having the same number of exons and matched internal junctions with the reference, while ISM includes isoforms with fewer 5’ exons but still matched internal junctions compared to the reference. The exact 5’ start and 3’ end can differ by any amount both for FSM and ISM. NIC includes isoforms without an FSM or ISM match but uses a combination of known donor or acceptor sites, whereas NNC refers to isoforms without FSM/ISM match and have at least one unannotated donor or acceptor site. Intergenic means the query isoform is in the intergenic region.

Figure 6C is a summary diagram presented to provide users with a quick and intuitive understanding of the software performance. Each method was scored based on its ranking in different evaluated aspects, with the top-ranked method receiving the highest score, and the score decreasing as the ranking drops. The score penalty is intensified if there is a significant difference in performance between two consecutive rankings, as observed in the case of TAMA’s memory and speed efficiency. Precision, sensitivity, speed, memory, scalability, and adaptability were all ranked based on the results of this benchmark study. Adaptability reflects how significant the method’s performance is influenced by changes in the five different factors investigated. Functional variability mainly demonstrates the diversity of tasks that the software is able to perform, with higher score awarded to software with more complex functions. For example, despite having only one additional functional module, FLAMES received the highest score for functional variability due to the complexity of its extra capacity, such as isoform detection from single cell TGS RNA-seq data. Usability is primarily assessed based on factors such as the smoothness of software installation, ease of usage (whether it is a one-line commander or requires additional scripting/contains multiple steps), and the frequency of encountering bugs during data processing.

## Data availability

The detailed information of the experimental datasets used in this study can be found in Table S1. The datasets used in this study are as follows: 1) Four TGS RNA-seq datasets from different types of human cells were collected from the NCBI with accession numbers SRR8568873, SRR8568871, SRR13762843, and SRR13762841^33, 34^, and the reference genome version used was GRCh38.105; 2) The ten mouse datasets can be found at NCBI with accession numbers SRR14630760, SRR14630758, SRR12800923, SRR12800924, SRR22522188, SRR22522033, SRR17960971, SRR17960979, SRR19257398, and SRR19257401^35–39^, and the reference genome used was GRCm39.105; 3) The NCBI accession numbers of the *Drosophila melanogaster* data include ERR3588905, and SRR13494726^40, 41^, and the reference genome used was Drosophila_melanogaster.BDGP6.32.53; 4) The five TGS direct RNA-seq datasets of *Caenorhabditis elegans* can be found with the NCBI accession numbers SRR8929006, SRR8929005, SRR8929004, SRR19055922, and SRR19055924^42, 43^; 5) The four newly generated naïve and primed hESCs TGS RNA-seq datasets can be accessed at NCBI’s Gene Expression Omnibus (GEO) via GEO Series accession number GSE227911 (https://www.ncbi.nlm.nih.gov/geo/query/acc.cgi?acc=GSE227911); 6) The NGS RNA-seq data of primed and naïve hESCs can be found from NCBI under the accession number SRR14073786, SRR14073787, SRR14073792, and SRR140737923^28^. Source data are provided with this paper.

## Code availability

Code for YASIM can be found on GitHub via https://github.com/WanluLiuLab/yasim/ or on PYPI https://pypi.org/project/yasim/. Documentation of YASIM can be found via https://labw.org/yasim-docs/. Code for profiler can be accessed via https://github.com/WanluLiuLab/labw_proc_profiler. Customized analysis code performed in this study can be found on GitHub via https://github.com/WanluLiuLab/2022_TGS_AS_Benchmark_Code.

## Acknowledgments

We express our gratitude to Dr. Aaron Trent Irving at ZJU-UoE Institute for help with the language editing. We would also like to acknowledge Dr. Chaochen Wang, and all lab members of the Liu lab at ZJU-UoE Institute for their helpful discussions. We also extend our thanks to the students of ZJU-UoE institute BMI-2019 class and staff from computational biology and system biology (2022-CBSB3) course in ZJU-UoE institute of Zhejiang University for their valuable suggestions on YASIM. We would also like to acknowledge the technical support provided by the Core Facilities of ZJU-UoE Institute. This work is supported by the Fundamental Research Funds for the Central Universities 226-2022-00134 (to W.L.), National Natural Science Foundation of China 32170551 (to W.L.), Alibaba Cloud (to W.L.), National Natural Science Foundation of China 32270835 (to D.C.), National Key Research and Development Program of China (No. 2022YFC2703503 to H.L.), and National Natural Science Foundation of China (No. 32250610202 to H.L.).

## Author contributions

Y.S. and W.L., conceived the study and designed experiments. Y.S., Z.Y., and W.L. wrote the manuscript. Z.Y., Y.S., R.Y., X.C., Z.X., and Y.G., implemented and compiled the documentation for YASIM. Z.Y. implemented the in-house profiler. D.C., H.L., and W.L. designed the hESCs experiments. S.J. and Z.A. performed the hESCs experiments. All authors contributed to the review and correction of the manuscripts.

## Competing interests

The authors declare no competing interest.

## Notes

### Competing Interest Statement

The authors have declared no competing interest.

